# Moderate nucleotide diversity in the Atlantic herring is associated with a low mutation rate

**DOI:** 10.1101/144303

**Authors:** Chungang Feng, Mats Pettersson, Sangeet Lamichhaney, Carl-Johan Rubin, Nima Rafati, Michele Casini, Arild Folkvord, Leif Andersson

## Abstract

The Atlantic herring (*Clupea harengus*) is one of the most abundant vertebrates on earth but its nucleotide diversity is moderate (π=0.3%), only three-fold higher than in human. The expected nucleotide diversity for selectively neutral alleles is a function of population size and the mutation rate, and it is strongly affected by demographic history. Here, we present a pedigree-based estimation of the mutation rate in the Atlantic herring. Based on whole-genome sequencing of four parents and 12 offspring, the estimated mutation rate is 1.7 × 10^−9^ per base per generation. There was no significant difference in the frequency of paternal and maternal mutations (8 and 7, respectively). Furthermore, we observed a high degree of parental mosaicism indicating that a large fraction of these *de novo* mutations occurred during early germ cell development when we do not expect a strong gender effect. The now estimated mutation rate – the lowest among vertebrates analyzed to date – partially explains the discrepancy between the rather low nucleotide diversity in herring and its huge census population size (>10^11^). But our analysis indicates that a species like the herring will never reach its expected nucleotide diversity for selectively neutral alleles primarily because of fluctuations in population size due to climate variation during the millions of years it takes to build up a high nucleotide diversity. In addition, background selection and selective sweeps lead to reductions in nucleotide diversity at linked neutral sites.

---

Empirical observations of nucleotide diversity in different species show that the variation is often much smaller than would be expected from simple population genetic models^1^. The Atlantic herring (*Clupea harengus*) is a good example of the paradox, since, in spite of an enormous census population size, its nucleotide diversity (π=0.3%)^2^ is middle-of-the-road when compared to terrestrial mammals, e.g. 0.1% for humans^3^ and 0.9% for European rabbits^4^ with much smaller census populations. A large census population does not necessarily mean that the effective population size (Ne) is large but the extremely low genetic differentiation at selectively neutral loci between geographically distant populations strongly suggests that current Ne must be high and genetic drift very low in the Atlantic herring^2^.

Before the NextGenerationSequencing-era, mutation rates were estimated by comparative genomics, by relating sequence differences to fossil record-dated estimates of species divergence times, or by tracking changes at specific loci in experimental studies. However, since species divergence is hard to date and the use of a small subset of loci can introduce bias, these methods have limited accuracy^5^. More recently, affordable whole genome sequencing has facilitated two approaches to estimate mutation rates: mutation accumulation lines and parent-offspring comparisons. The mutation accumulation approach, where an inbred line is maintained for a number of generations and the mutation rate is measured by counting up differences between the first and last generation, has the advantage of scalability, since it is possible to increase the number of mutation events observed by including more generations. On the other hand, the approach requires an organism that can be reproduced as viable inbred lines, and it is difficult to fully eliminate purifying selection against deleterious new mutations. The parent-offspring approach, which relies on using high coverage whole-genome sequencing to detect differences between parents and their offspring, alleviates the cultivation related issues, and has thus become the preferred method for estimating the mutation rate in non-model organisms. The trade-off is that the total number of mutation events per progeny will typically be small.

Currently, the number of studies using any of the methods outlined above remains small, and the available data is somewhat biased towards unicellular organisms^6–11^, insects^12–14^ and mammals^15–18^, while including a single plant^19^ and one bird^20^. In all, this leaves large sections of the tree of life essentially unexplored. This is problematic for drawing general conclusions about the relationship between neutral diversity, effective population size and mutation rate, which is a topic of considerable interest in population genetics^1,21^.

In this study, to our knowledge the first of its kind in a teleost, we estimate the genome-wide point mutation rate in Atlantic herring. The Atlantic herring was chosen due to its suitability as a population genetic model system; it is one of the most abundant vertebrate species on earth with external reproduction involving large numbers of gametes per reproducing adult. In essence, these properties make the Atlantic herring one of the best approximations of a randomly mating, infinite size population among vertebrates. In addition, there exists a high-quality draft genome assembly^2^, which is a pre-requisite for a study of this kind. We have employed the parent-offspring approach, and base our measurement on two families, each containing two parents and six offspring. We here estimate the spontaneous mutation rate to be 1.7 × 10^−9^ per site per generation in the Atlantic herring, eight-fold lower than the rate in humans and the lowest rate reported so far for a vertebrate.

## Results

### Whole genome sequencing and variant calling

We have generated two-generation experimental pedigrees for spring-spawning Atlantic and Baltic herring (classified as a subspecies of the Atlantic herring by Linneaus^22^), each comprising the two parents and six offspring (Table 1). We performed whole-genome sequencing of these two pedigrees using genomic DNA isolated from muscle tissue. As detection of *de novo* mutations requires high sequence coverage, we sequenced each individual to ~ 45-71x (Table 1), in line with the procedures used in previous studies^13,16^. The sequences were aligned to the recently published Atlantic herring genome^2^. A total of 5.3 (Atlantic) and 5.2 (Baltic) million raw SNPs were detected in each pedigree, respectively, using GATK (see Methods)^23^.

**Table 1.** Summary of the pedigrees used for whole-genome sequencing

| No | ID | Pedigree | Sequencing<br>depth (x) | *de novo*<br>mutations |
| --- | --- | --- | --- | --- |
| <u>Pedigree 1, Atlantic herring</u> |  |  |  |  |
| 1 | AM8 | Father | 65.7 | N.A. |
| 2 | AF8 | Mother | 70.2 | N.A. |
| 3 | AA1 | Offspring | 65.6 | 1 |
| 4 | AA2 | Offspring | 70.9 | 2 |
| 5 | AA3 | Offspring | 47.2 | 0 |
| 6 | AA4 | Offspring | 66.9 | 3 |
| 7 | AA5 | Offspring | 64.2 | 4 |
| 8 | AA6 | Offspring | 61.2 | 1 |
| <u>Pedigree 2, Baltic herring</u> |  |  |  |  |
| 9 | BM19 | Father | 71.8 | N.A. |
| 10 | BF21 | Mother | 65.1 | N.A. |
| 11 | BB1 | Offspring | 74.5 | 2 |
| 12 | BB2 | Offspring | 61.6 | 1 |
| 13 | BB3 | Offspring | 75.0 | 0 |
| 14 | BB4 | Offspring | 69.9 | 2 |
| 15 | BB5 | Offspring | 60.6 | 2 |
| 16 | BB6 | Offspring | 62.6 | 1 |

N.A. = Not available.

### *Identification and validation of the* de novo *mutations*

Detection of *de novo* mutations with high confidence requires a careful examination of raw variant calls and application of highly stringent filtering criteria. Using a standard genotype-calling pipeline will typically lead to the great majority of novel sequence variants detected being false positives. Screening of provisional candidate mutations in a single offspring indicated that this was the case, as many candidates could not be verified using Sanger sequencing. Hence, in order to minimize the frequency of false positives by the *de novo* calls using only the GATK variant caller, we separately performed variant calling using SAMTOOLS^24^ and only selected novel mutations detected by both variant callers (Figure 1). In addition, we applied strict filtering criteria in order to remove variants detected due to sequencing and alignment errors. We excluded variant calls from genomic regions with low mappability (see Methods) and repetitive regions detected by Repeat Masker^25^. Furthermore, we defined the cut-off parameters for sequence depth, SNP and genotype quality-related statistics using the set of SNPs that were fixed for different alleles in both parents and thus heterozygous in all offspring (Figure 1, Methods). As this strict filtering could lead to failure to detect some fraction of true heterozygotes, we estimated the false negative rate of our pipeline by calling SNPs in each individual offspring separately, in order to eliminate bias stemming from shared SNPs present in multiple individuals being called with higher power. For this analysis we used 116,910 polymorphic sites where the parents were homozygous for different alleles in the joint genotype calling.

The expectation is that these sites are heterozygous in all offspring, but that information did not influence SNP calling. By separating the individuals, we mimicked the situation for *de novo* mutations, which are typically not shared. Using the same pipeline as for the *de novo* detection, the average detection rate of such heterozygous positions across all offspring was 94.1%, yielding a false negative rate of 5.9%. As an alternative way of estimating the false negative rate, we used a simulation procedure where we generated mutated reads for 1,000 positions within callable regions. Each site in each offspring had its frequency of mutated reads determined by a sample from the observed frequency distribution of called heterozygous sites in the original data set; see *Methods* for details. Across all offspring, we found an overall frequency of 2.7% false negative calls, while roughly 9% of sites failed to generate a call (Supplementary File 1). Overall, the two methods used are in agreement. However, for the purpose of the final calculations we will use the empirical estimate of 5.9%, which includes both incorrect and failed calls, as it is derived directly from the real data set. The choice have minor effects on the estimated mutation rate, as using the simulated value would result in the finale rate being approximately 5% higher.

This stringent filtering procedure identified a total of 17 candidate *de novo* mutations, nine in the Atlantic pedigree and eight in the Baltic one (Tables 1 and 2). Two of the 17 *de novo* mutations were each found in two different offspring from the same pedigree.

We performed Sanger sequencing of the genomic regions around each of these putative *de novo* mutations in all parents and the 12 offspring (Figure 1—figure supplement 1). This confirmed that all 17 putative *de novo* mutation events were genuine and all the peak ratios of two alleles were close to 1:1 consistent with germ-line mutations. This means that the observed false positive rate across all candidate sites was zero.

In order to estimate the transmission frequencies of our detected *de novo* mutations, we measured the rate of transfer of the *de novo* mutations in a larger set of offspring (*n* = 46 and 50 per family), in order to infer when during the formation of the parental germ line the mutation occurred (Table 2). For eight out of seventeen *de novo* mutations we observed more than one sibling carrying exactly the same mutation (Table 2). The range of occurrences for the *de novo* mutations was one to nine among the 50 offspring. Even the maximum of the observed transfer rates (18% for scaffold.153: 2,684,380 T>G) was significantly lower than the 50% expected for a fixed mutation (*P* = 1.4 × 10^−3^, Fisher’s exact test). About half of the *de novo* mutations were present in two or more offspring, indicating that they occurred during early germ cell divisions. Assuming that the number of cell divisions from zygote to mature sperm or egg is similar in Atlantic herring to the one in mammalian species, we can conclude from a recent simulation study^15^ that it would be highly unlikely to observe such a high rate of parental mosaicism unless a large fraction of the *de novo* mutations occurred during early germ cell divisions. Further, we detected a higher incidence of parental mosaicism in the Atlantic herring than in the Baltic herring pedigree (Table 2; *P* = 0.01, Fisher’s exact test). The finding that the same mutation was observed in two or more siblings for eight of the putative *de novo* mutations confirms that these must be germ-line mutations and not somatic mutations.

**Table 2.** Summary of the *de novo* mutations identified in Atlantic herring.

| SNP position<br>scaffold:position | Id | Mutation |  | Freq<sup>†</sup> | Origin<sup>††</sup> | Type<sup>†††</sup> | Region |
| --- | --- | --- | --- | --- | --- | --- | --- |
|  |  | Ref | Var |  |  |  |  |
| 1157:174,127 | AA4 | T | A | 1/50 (-) | M | TV | Intergenic |
| 153:2,684,380 | AA2 | T | G | 9/50 (18%) | P | TV | Intronic |
| 241:7,752,158 | AA5 | C | A | 5/50 (10%) | M | TV | Intergenic |
| 4:5,098,858 | AA5 | T | C | 2/50 (4%) | M | TS | Intronic |
| 481:1,927,799 | AA4, AA5* | C | A | 6/50 (12%) | P | TV | 3' UTR |
| 61:815,077 | AA4 | A | T | 3/50 (6%) | N.A. | TV | Intergenic |
| 62:613,919 | AA1, AA6* | C | A | 6/50 (12%) | M | TV | Intergenic |
| 729:1,499,224 | AA2 | C | T | 4/50 (8%) | M | TS | Intronic |
| 887:195,946 | AA5 | G | A | 1/50 (-) | P | TS | Intronic |
| 10:1,443,002 | BB4 | C | T | 1/46 (-) | P | TS | Intronic |
| 151:267,875 | BB5 | A | T | 1/46 (-) | P | TV | Exonic |
| 177:1,045,894 | BB1 | A | G | 1/46 (-) | P | TS | Intronic |
| 194:478,776 | BB6 | A | G | 1/46 (-) | N.A. | TS | Intronic |
| 246:1,890,479 | BB4 | T | C | 1/46 (-) | P | TS | Intergenic |
| 257:380,993 | BB2 | G | A | 1/46 (-) | M | TS | Intergenic |
| 26:2,976,192 | BB1 | T | C | 2/46 (4%) | P | TS | Intronic |
| 37:1,374,669 | BB5 | G | A | 1/46 (-) | M | TS | Intronic |

\*Same mutation detected in two progeny.
<sup>†</sup>Number of siblings carrying the *de novo* mutation; - the frequency of transmission was only estimated when two or more progeny with the *de novo* mutation was detected.
<sup>††</sup>M:Maternal, P:Paternal, N.A. = Not available
<sup>†††</sup>TV = Transversion, TS = Transition

### Parental origin of de novo mutations

We also explored if the 17 germ line *de novo* mutations had paternal or maternal origin. For 14 of the *de novo* mutations, we could detect an additional segregating site within the same Illumina sequencing read (length 125 bp) or mate-pair read that spanned the respective *de novo* mutation and was uniquely associated with either parent. In these cases, the parental origin could be directly inferred. We were able to infer the parental origins of one additional *de novo* mutation by PCR cloning and sequencing.

Out of the 15 mutations for which their parental origin was determined, there was no significant difference between paternal (eight) and maternal (seven) mutations (Table 2). Paternal bias in the origin of *de novo* mutations has been shown in mammals, such as human (ratio = 3.9)^16^ and chimpanzee (ratio = 5.5)^18^, where the main reason is thought to be the larger number of cell divisions during spermatogenesis than during oogenesis^26^. While the numbers are small, a binomial test against the human ratio indicates that the gender bias in herring, if it exists at all, is significantly weaker than in humans (*P* = 0.004). In herring, both sexes produce large numbers of gametes and males only produce sperms during the spawning season (a few months per year). Furthermore, the high degree of parental mosaicism indicates that a large fraction of the *de novo* mutations reported here must have occurred during early germ cell development when we do not expect a strong gender effect. These circumstances offer a reasonable explanation to the balanced parental origin of *de novo* mutations in the Atlantic herring.

### Characteristics of de novo mutations

Among the 17 *de novo* mutations, there were 10 transitions and seven transversions, yielding a transition/transversion ratio of 1.4. An overrepresentation of transitions is expected, and the observed ratio falls in the range found in previous *de novo* mutation studies. For example, Kong *et al*. identified 3,344 transitions out of 4,933 events (ratio = 2.1) in humans^16^, while Keightley *et al*. found five out of nine events (ratio = 1.25) in the tropical butterfly *Heliconius melpomene*^1^. In humans and other mammals there is a well-established excess of CpG>TpG mutations^16^. There was no such strong trend in our small dataset as only 1 out of 17 *de novo* mutations was of this type.

There were six mutations located in intergenic regions, nine intronic mutations, one 3’ UTR mutation and one exonic mutation. In all, this is a distribution that does not deviate significantly from random expectation, given the composition of the genome after mappability filtering (*P* = 0.65, Fisher’s exact test).

### Estimation of mutation rates

We identified nine and eight *de novo* mutations in the Atlantic herring and the Baltic herring pedigrees, respectively. Since we had 12 progeny in total, our estimate of the number of events per meiosis is 0.71 (17/24). After strict filtering of genomic regions with low mappability and repetitive sequences, we had ~ 442 Mb of sequence available for variant screening (representing ~ 52% of the genome). Based on the distribution of read coverage in a random subset of the genome (Supplementary File 1), we expect 2.6% of this region to have insufficient depth for a successful SNP call, giving us a final calla ble region of 442 × 0.974 = 431 Mb. The mutation rate per site per generation can thus be estimated as 17/(2 × 12 × 431 × 10^6^) = 1.6 × 10^−9^ (95% CI = 0.9 − 2.5 × 10^−9^, assuming that the mutations are Poisson distributed). If we correct for the estimated false negative rate (5.9%) we obtain nearly identical numbers: 1.7 × 10^−9^ (95% CI = 0.9 − 2.7 × 10^−9^). It should be noted that this number reflects the rate in the callable fraction of the genome, which by definition does not contain repeat regions. Thus, the true genomic average could be somewhat higher, as replication of repetitive regions tends to be more error-prone, but the decreased calling power in those regions makes diversity hard to estimate in an unbiased fashion. However, these issues are not unique to the Atlantic herring, similar caveats apply to estimates of mutation rates in other species as well, and the results should thus be comparable across species.

Based on historical sampling of several herring stocks, we estimated the minimum generation time of Atlantic herring before the onset of large-scale commercial fishing to be approximately six years (Supplementary file 2). Using this historical generation time, the mutation rate per site per year in the Atlantic herring was estimated at 2.9 × 10^−10^ (95% CI = 1.5 × 10^−10^ − 4.5 × 10^−10^).

## Discussion

This study provides new insights regarding factors affecting the mutation rate and levels of nucleotide diversity in vertebrates. Our finding of a high degree of parental mosaicism for the detected *de novo* mutations is consistent with a recent study indicating that the early cleavage cell divisions in the germ-line are particularly mutation-prone^15^. A high rate of *de novo* mutations at early germ-cell divisions has also been reported for *Drosophila*^27^.

The now estimated mutation rate (1.7 × 10^−9^) is on the low end compared to what would be expected from the previously established empirical trends between genome size, effective population size, and mutation rate among eukaryotes^21^. In fact, the estimated mutation rate (μ) for the Atlantic herring is the lowest for a vertebrate species to date (Table 3); about eight-fold lower than in humans. By combining this value with the neutral diversity level (π=0.003) found by Martinez Barrio *et al*.^2^ and the expected relationship between nucleotide diversity, the mutation rate and effective population size (N_e_) for selectively neutral alleles (π =4N_e_ μ), we obtain an estimated N_e_ of approximately 5 × 10^5^. While this number is larger than for most terrestrial animal species, it is still much lower than the census population size of the herring, which must be on the order of 10^11^ or higher. There are several factors that may contribute to this discrepancy, with demographic history being a clear candidate. Using coalescent analysis and allele frequency distributions, Martinez Barrio *et al*.^2^ showed that the herring population is expanding from a relatively strong bottleneck during the most recent glaciation period (starting approximately 2 MYA). Since the diversity-based estimate of effective population size can be considered as an average over time this bottleneck still have a major impact on the current nucleotide diversity. Population genetics theory implies that it will take 4Ne generations before populations reach their genetic equilibrium^28^. We have estimated the generation interval to approximately six years in this study (Supplementary file 2) and a conservative estimate of the current (not long-term) N_e_ is 10^7^, which appears reasonable since we estimated long-term Ne at 5 × 10^5^ and we have evidence for population expansion (e.g. excess of rare alleles^2^). These figures indicate that it will take about 240 million years before the herring populations reach genetic equilibrium! Thus, it is obvious that a species with a huge population size like the herring and a relatively long generation interval will never reach genetic equilibrium. Background selection (the elimination of deleterious alleles) and selective sweeps will also lead to reductions in nucleotide diversity at linked neutral sites. Furthermore, highly efficient purifying selection decreases the fraction of the genome that is selectively neutral which is also expected to lead to a slightly reduced nucleotide diversity.

**Table 3.**
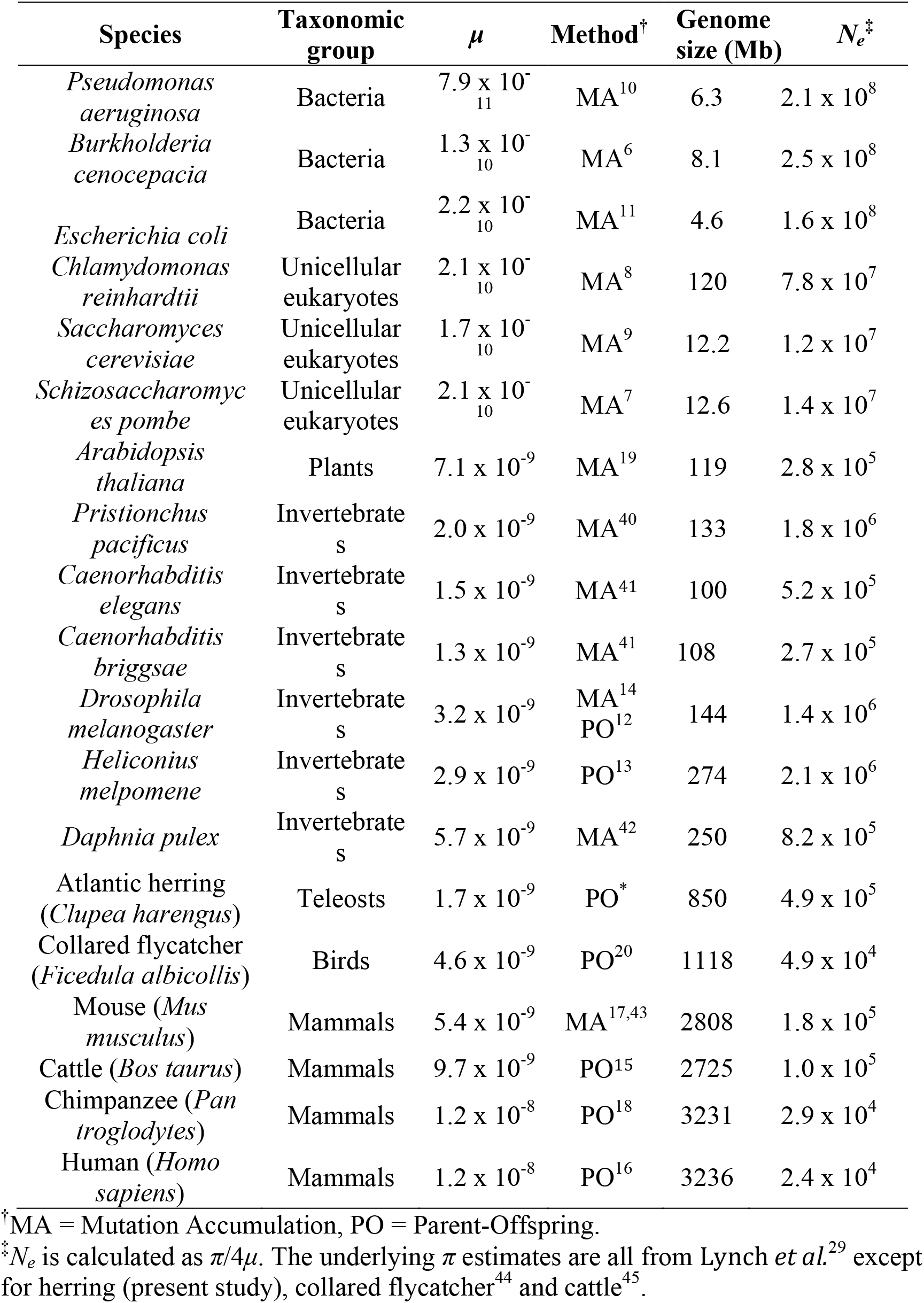
Summary of mutation rates measured to date.

| Species | Taxonomic group | $\mu$ | Method<sup>†</sup> | Genome size (Mb) | $N_e^{\ddagger}$ |
| --- | --- | --- | --- | --- | --- |
| *Pseudomonas aeruginosa* | Bacteria | $7.9 \times 10^{-11}$ | MA<sup>10</sup> | 6.3 | $2.1 \times 10^8$ |
| *Burkholderia cenocepacia* | Bacteria | $1.3 \times 10^{-10}$ | MA<sup>6</sup> | 8.1 | $2.5 \times 10^8$ |
| *Escherichia coli* | Bacteria | $2.2 \times 10^{-10}$ | MA<sup>11</sup> | 4.6 | $1.6 \times 10^8$ |
| *Chlamydomonas reinhardtii* | Unicellular eukaryotes | $2.1 \times 10^{-10}$ | MA<sup>8</sup> | 120 | $7.8 \times 10^7$ |
| *Saccharomyces cerevisiae* | Unicellular eukaryotes | $1.7 \times 10^{-10}$ | MA<sup>9</sup> | 12.2 | $1.2 \times 10^7$ |
| *Schizosaccharomyces pombe* | Unicellular eukaryotes | $2.1 \times 10^{-10}$ | MA<sup>7</sup> | 12.6 | $1.4 \times 10^7$ |
| *Arabidopsis thaliana* | Plants | $7.1 \times 10^{-9}$ | MA<sup>19</sup> | 119 | $2.8 \times 10^5$ |
| *Pristionchus pacificus* | Invertebrates | $2.0 \times 10^{-9}$ | MA<sup>40</sup> | 133 | $1.8 \times 10^6$ |
| *Caenorhabditis elegans* | Invertebrates | $1.5 \times 10^{-9}$ | MA<sup>41</sup> | 100 | $5.2 \times 10^5$ |
| *Caenorhabditis briggsae* | Invertebrates | $1.3 \times 10^{-9}$ | MA<sup>41</sup> | 108 | $2.7 \times 10^5$ |
| *Drosophila melanogaster* | Invertebrates | $3.2 \times 10^{-9}$ | MA<sup>14</sup><br>PO<sup>12</sup> | 144 | $1.4 \times 10^6$ |
| *Heliconius melpomene* | Invertebrates | $2.9 \times 10^{-9}$ | PO<sup>13</sup> | 274 | $2.1 \times 10^6$ |
| *Daphnia pulex* | Invertebrates | $5.7 \times 10^{-9}$ | MA<sup>42</sup> | 250 | $8.2 \times 10^5$ |
| Atlantic herring (*Clupea harengus*) | Teleosts | $1.7 \times 10^{-9}$ | PO<sup>*</sup> | 850 | $4.9 \times 10^5$ |
| Collared flycatcher (*Ficedula albicollis*) | Birds | $4.6 \times 10^{-9}$ | PO<sup>20</sup> | 1118 | $4.9 \times 10^4$ |
| Mouse (*Mus musculus*) | Mammals | $5.4 \times 10^{-9}$ | MA<sup>17,43</sup> | 2808 | $1.8 \times 10^5$ |
| Cattle (*Bos taurus*) | Mammals | $9.7 \times 10^{-9}$ | PO<sup>15</sup> | 2725 | $1.0 \times 10^5$ |
| Chimpanzee (*Pan troglodytes*) | Mammals | $1.2 \times 10^{-8}$ | PO<sup>18</sup> | 3231 | $2.9 \times 10^4$ |
| Human (*Homo sapiens*) | Mammals | $1.2 \times 10^{-8}$ | PO<sup>16</sup> | 3236 | $2.4 \times 10^4$ |

<sup>†</sup>MA = Mutation Accumulation, PO = Parent-Offspring.
<sup>‡</sup> $N_e$ is calculated as $\pi/4\mu$ . The underlying $\pi$ estimates are all from Lynch *et al.*<sup>29</sup> except for herring (present study), collared flycatcher<sup>44</sup> and cattle<sup>45</sup>.

The fact that the observed mutation rate is unusually low in the Atlantic herring is of interest in relation to the drift-barrier hypothesis^29^, which predicts that the purging of slightly deleterious mutations affecting the mutation rate is particularly effective in species that have a very large population size, large fecundity and close to random mating, conditions which the Atlantic herring meets (Table 3). However, since the population size of the Atlantic herring appears to have fluctuated over time^2^, it remains unclear exactly how powerful selection has been in a time-average perspective, which means the support for the drift-barrier hypothesis is not unconditional. Additionally, the low body temperature of a marine fish may also slow down the metabolic rate which has been suggested to decrease the mutation rate^30^, there is thus a need to compare with mutation rates from other species, with lower populations sizes but similar body temperatures, before we can draw firm conclusions about the relationship between population size and mutation rate.

According to simple, ideal-case population genetic models there should be a positive relationship between nucleotide diversity and population size, so that a population at mutation-drift balance has a nucleotide diversity of 4*N_e_μ*. However, as outlined above, this expectation is disrupted by population size fluctuations over time and selective forces. In practice, population sizes are only weakly, if at all, correlated with nucleotide diversity^1^. Our finding that the inherent mutation rate is approximately eight times lower in Atlantic herring than in humans indicates that differences in intrinsic mutation rate is also an important factor when comparing nucleotide diversities among species. In the case of the Atlantic herring, the low mutation rate, in conjunction with demographic history and efficient purifying selection, explains the majority of the apparent disparity between nucleotide diversity and the census population size in the Atlantic herring.

## Methods

### Sample

Two full-sib families were generated by crossing wild-caught Atlantic herring from Bergen (Norway) and Baltic herring from Hästskär (Sweden). For each family, six offspring from a total of 50 progeny were selected for sequencing together with the two parents. Our aim was to determine the mutation rate to its order of magnitude and one to two significant digits. Thus, a samples size of 12 progeny was expected to result in about 100 detectable novel mutations based on previously known vertebrate mutation rates and the size of the genomic regions we could use to detect mutations. Genomic DNA was isolated from muscle tissue using Qiagen DNeasy Blood and Tissue kit. DNA libraries were constructed using the TruSeq PCR-free kit.

### Whole-genome sequencing

All individuals were sequenced on Illumina HiSeq2500 machines, using 2 × 125 bp paired reads to a sequencing depth of ~47–71X. The short reads were aligned to the *Clupea harengus* reference genome (v1.2)^2^ using BWA v0.6.2^31^ with default parameters. The data were then filtered based on mappability, calculated using GEM^32^, within the reference assembly, so that only positions with mappability 1 that were also inside 1 kb windows with average mappability > 0.95 were included in the downstream analysis; 442 Mb (52%) of genome sequence passed this filtering step. The sequence data have been deposited in the SRA archive (PRJNA356817).

### Variant calling and filtration

Sequence alignments from the previous step were used for calling variants using two separate tools; GATK v3.3.0^23^ and SAMTOOLS v.1.19^24^. We used GATK HaplotypeCaller with default parameters that performs simultaneous calling of SNP and Indels via local *de novo* assembly of haplotypes (see GATK manual for details). We ran HaplotypeCaller separately for each individual to generate intermediate genomic VCF^33^ files (gVCF). Afterwards, we used the GenotypeGVCFs module in GATK to merge gVCF records from each individual (altogether 12 from the two pedigrees) using the multi-sample joint aggregation step that combines all records, generate correct genotype likelihood, re-genotype the newly merged record and re-annotate each of the called variants and thereby generate a VCF file. For SAMTOOLS, we used the standard multi-sample SNP calling pipeline^24^ using mpileup module for calling raw variants.

Once we got the raw variant calls, we filtered small insertions and deletions and only used SNPs for downstream analysis. Furthermore, we also removed SNPs that had missing genotypes in one or both parents, as these SNPs were not informative. Afterwards, we extracted a subset of SNPs where parents were homozygous for different allele and all six offspring were heterozygous (the genotype calls were considered heterozygous in offspring if the minor allele frequency was > 25%). The SNP quality annotations in this set of “known” heterozygous offspring were used as proxy to consider the quality parameter of true SNPs in the dataset. We extracted various SNP quality annotations recorded in the VCF file like total read depth, mapping quality, mapping quality rank sum, base quality, base quality rank sum, read position rank sum, quality by depth, genotype quality, allele depth (see GATK manual for details on these parameters) and examined their distributions in the subset of our known heterozygous offspring. As the distributions of these quality parameters were close to being normal distribution (data not shown), we used the threshold of mean ±2 × standard deviation for each of these quality estimates as the standard cut-offs for our in-house SNP filtering pipeline to filter raw SNPs in our entire dataset (Figure 1).

### De novo mutation calling

From the filtered SNP dataset generated in the previous step, we further selected those sites where both parents were homozyogous for the reference allele and at least one offspring carried the variant allele in the heterozygous or homozygous state. These two sets of raw novel mutations in offspring independently called by GATK and SAMTOOLS were then intersected and the sites that were detected by both variant callers were considered as our true *de novo* mutations among the progeny.

### Experimental validation and parental origin

PCR amplification and Sanger sequencing of both strands verified all candidate mutations. We inferred the parental origin of the *de novo* mutations based on flanking SNP alleles that could be verified by Sanger sequencing and only have been transferred from one of the parents. The parental origin of fifteen *de novo* mutations could be directly deduced from SNP alleles segregating between the two parents present on the same short Illumina read and mate-pair read as the *de novo* mutation (at least 5 reads). The other *de novo* mutations were determined via cloning PCR fragments and sequencing; we sequenced at least 7 independent clones for each *de novo* mutation.

### Estimation of the false negative rate

Firstly, we estimated the false negative rate by performing genotype calls at those nucleotide positions where the parents were fixed for different alleles. The genotype calls for progeny were done without using the information for parents to mimic the detection of *de novo* mutations. Secondly, we also used simulation to estimate the false negative rate. We selected regions with mappability of 1 without any polymorphism. From these regions, we selected approximately 1,000 sites for each offspring and then introduced *de novo* mutations. Then, we aligned the new reads and called SNPs using the pipeline described in Figure 1. Finally, we compared the SNP calls with expected genotypes based on the mutated sites and calculated the false negative rate.

### Estimation of generation time

The generation length of populations with overlapping generations is equal to the mean age of parents^34^. Following Miller and Kapuscinski^35^, this was approximated as the mean age of spawners (age-specific number of fish multiplied by the age-specific proportion of reproductive fish) weighted by age-specific mean weights. In our analyses we used age-specific weights as proxy for age-specific fecundity, since in Atlantic herring weights and fecundity are strongly and nearly linearly correlated^36,37^. We estimated the generation time for the herring stocks with data starting shortly after the end of the World War II, a period characterized by still low commercial exploitation which started to increase after the early 1960s. The stocks were the North Sea/Skagerrak/Kattegat/English Channel, the Celtic Sea, the West of Scotland/West of Ireland, the Irish Sea and the Norwegian spring spawning herring. Data on age-specific abundance, maturity and mean weight were extracted from stock assessment reports^38,39^.

The generation time was very similar for almost all the stocks in the first available period after the World War II, characterized by low exploitation, i.e. in 1947-1965. During this period, the generation time declined between ~6 years in late 1940s (corresponding to the lowest exploitation) and ~5 years in 1965, decreasing further in successive years. No data were available for the period before 1947 when the generation time was likely to have been higher. The Norwegian spring spawning herring showed a higher generation time than the other stocks, oscillating around 10 years in the 1950s. We therefore consider the generation time of 6 years as a minimum estimate for Atlantic herring under no or moderate exploitation.

## Acknowledgements

We thank Michel Georges for comments on the manuscript. The work was funded by the ERC project BATESON (to LA), the GENSINC project funded by the Norwegian Research Council (to AF and LA). Illumina sequencing was performed by the SNP&SEQ Technology Platform, supported by Uppsala University and Hospital, SciLifeLab and Swedish Research Council (80576801 and 70374401). Computer resources were provided by UPPMAX, Uppsala University.

## Author contributions

LA conceived the study. LA, CF, MP, SL designed the experiment. CF performed the experimental work. MP and SL carried out the bioinformatic analysis. C-JR and NR contributed to the bioinformatic analysis. MC estimated the generation time in Atlantic herring. AF was responsible for generating the pedigree material. LA, CF, MP, SL wrote the paper with contributions from all authors.

## Figure legends

**Figure. 1: Flowchart describing the *de novo* mutation-calling pipeline.** A schematic illustration of the steps used in calling and filtering the candidate mutations.

**Figure 1—figure supplement 1. Sanger sequencing chromatograms of the *de novo* mutations.** Chromatograms from the identified target offspring and its parents for each region containing a candidate *de novo* mutation.

---

**Supplementary File 1:** Summary statistics of the SNP calls underlying the estimation of the false negative rate by means of simulation.

**Supplementary File 2:** Estimates of generation time for different stocks of herring.

